# Partitioning the roles of insect and microbial enzymes in the metabolism of the insecticide imidacloprid in *Drosophila melanogaster*

**DOI:** 10.1101/134122

**Authors:** Roberto Fusetto, Shane Denecke, Trent Perry, Richard A. J. O’Hair, Philip Batterham

## Abstract

Resistance to insecticides through enhanced metabolism is a worldwide problem. The *Cyp6g1* gene of the vinegar fly, *Drosophila melanogaster*, is a paradigm for the study of metabolic resistance. Constitutive overexpression of this gene confers resistance to several chemical classes of insecticides, including the neonicotinoids exemplified by the insecticide imidacloprid (IMI). The metabolism of IMI in this species has been previously shown to yield oxidative and nitro-reduced metabolites. While levels of the oxidative metabolites are correlated with CYP6G1 expression, nitro-reduced metabolites are not, raising the question of how these metabolites are produced. Some IMI metabolites are known to be toxic, making their fate within the insect a second question of interest. These questions have been addressed by coupling the genetic tools of gene overexpression and CRISPR gene knock-out with the sensitive mass spectrometric technique, the Twin-Ion Method (TIM). Analysing axenic larvae indicated that microbes living within *D. melanogaster* are largely responsible for the production of the nitro-reduced metabolites. Knock-out of *Cyp6g1* revealed functional redundancy, with some metabolites produced by CYP6G1 still detected. IMI metabolism was shown to produce toxic products that are not further metabolized but readily excreted, even when produced in the Central Nervous System (CNS), highlighting the significance of transport and excretion in metabolic resistance.

## Introduction

Insecticides have long been used to control insect pests that negatively impact agriculture and human health. Tobacco extracts and then pure (S)-nicotine, targeting nicotinic acetylcholine receptors (nAChRs), were the first insecticides used against pests around the world^1^. While effective, (S)-nicotine is more toxic to mammals than insects so advances in organic synthesis were exploited to produce nicotine derivatives (neonicotinoids), notably IMI, that have a higher affinity for insect nAChRs^2-4^. Recently, there have been major concerns about the impact of honeybee exposure to IMI and some other neonicotinoids^5-7^ and the increasing number of pest species evolving resistance^8-10^. Determining the mechanisms by which pests develop resistance to insecticides is crucial to develop new control strategies for insecticide resistant pests. The vinegar fly, *Drosophila melanogaster*, has been shown to be a useful model system for the study of resistance mechanisms^11,12^.

The most common mechanism of neonicotinoid resistance involves the constitutive overexpression of drug metabolising enzymes (DMEs), particularly cytochrome P450 monooxygenases of the CYP6 family^13-15^. *Cyp6g1* from *D. melanogaster* is the best characterized gene of this family^16^. Resistance to insecticides of diverse chemical structure, including IMI, is due to elevated transcript levels in key metabolic tissues (midgut, fat body and malpighian tubules), caused by the insertion of a long terminal repeat of the *Accord* retrotransposon upstream of the gene^16-18^. Further transposable element insertions and gene duplications have served to increase the levels of *Cyp6g1* transcript and resistance in natural populations around the world^19,20^. In the laboratory, it has been possible to reproduce such resistance using controlled tissue specific gene expression with the GAL4/UAS system and an *Accord* driver^17,21^.

Two distinct pathways of IMI metabolism have been observed in nature: oxidation and nitro-reduction (Figure 1). While both mechanisms have been established in mammals^22,23^ and plants^24,25^, nitro-reduction of IMI is the main mechanism used by soil bacteria^26-28^ with the only exception being *Stenotrophomonas maltophilia*, which predominantly produces IMI-5-OH^29^. In insects, IMI-5-OH and IMI-Ole are the key metabolites identified in different insect species^30,31^. However, the predominant identification of IMI-Urea and 6-chloronicotinic acid (6-CNA) in midgut and rectum tissues of *Apis mellifera*^32^ and the detection of nitro-reduced metabolites as major metabolites in the eastern subterranean termite, *Reticulitermes flavipes*^33^, show that IMI metabolism in insects is far from being understood.

**Figure 1.**
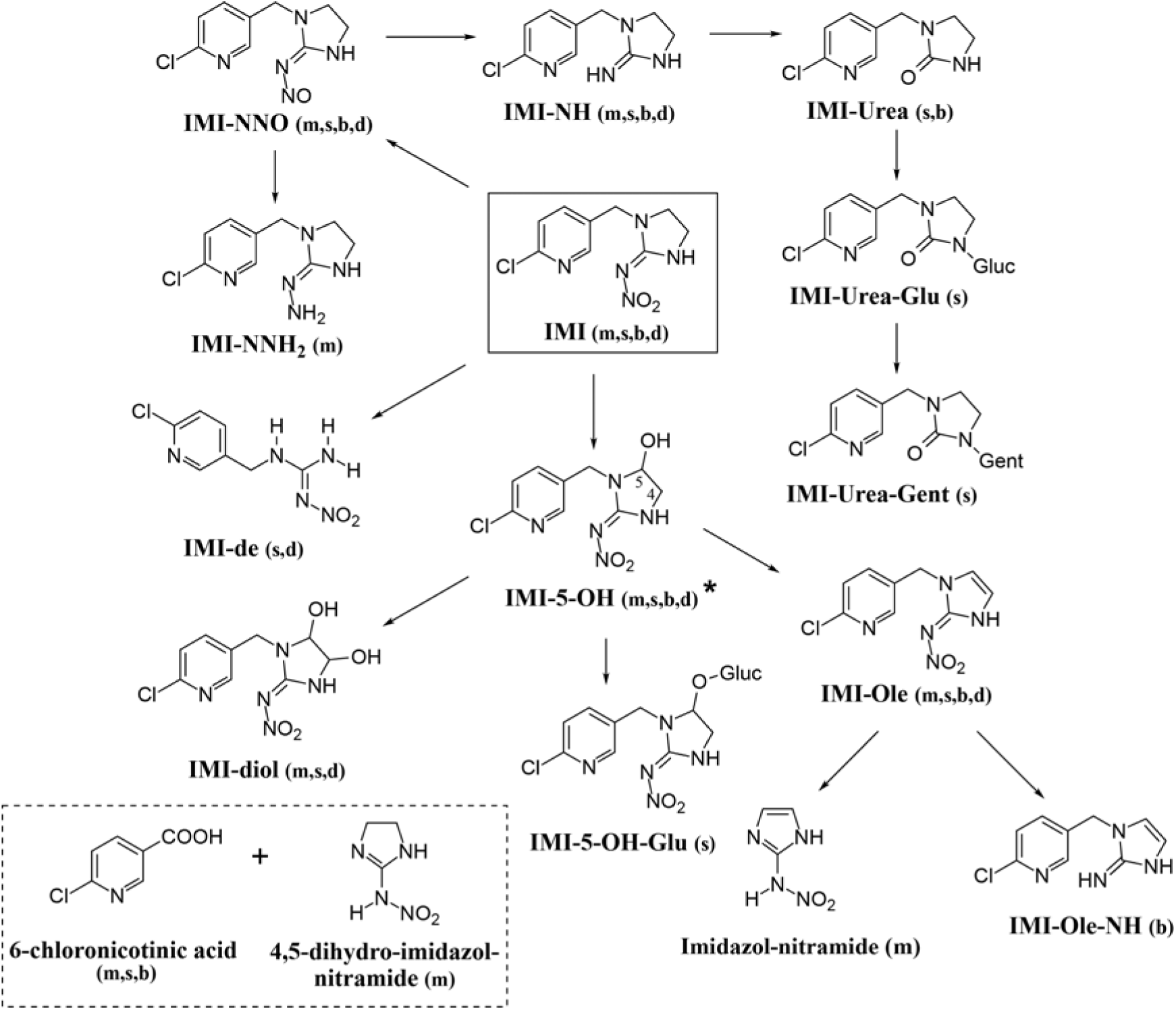
The metabolites of IMI identified in mice^22^ (m), spinach^24^ (s), soil bacteria^28^ (b), and *D. melanogaster*^35^ (d). Two main mechanisms of metabolism: oxidation and nitro-reduction. Oxidation on the imidazolidine ring of IMI produces IMI-5-OH and IMI-diol, while nitro-reduction generates IMI-NNO, IMI-NNH_2_, IMI-NH and IMI-Urea. Other metabolic reactions observed are dehydration (IMI-Ole), O-and N-conjugation of glucose (IMI-5-OH-Glu and IMI-Urea-Glu) and gentamicin (IMI-Urea-Gent) and deethylation of IMI (IMI-de). Formation of 6-Chloronicotinic acid and 4,5-dihydro-imidzol-nitramide is observed as direct products of IMI (dashed box). (*) Hydroxylation of IMI in position 4 on the imidazolidine ring, IMI-4-OH, was reported in *Nicotiana tabacum* cell cultures expressing CYP6G1^34^.

The role of CYP6G1 in IMI metabolism was first examined using heterologous expression in tobacco (*Nicotiana tabacum*) cell culture^34^. Subsequently, the integration of the TIM with the selective tissue expression of *Cyp6g1* using the GAL4/UAS system permitted the *in vivo* study of IMI metabolism in *D. melanogaster*^35^. The oxidation products IMI-5-OH, IMI-diol, IMI-Ole and IMI-de were detected at levels that correlated with *Cyp6g1* expression in the insect. Significantly, nitro-reduced derivatives, IMI-NNO and IMI-NH, were also detected but their levels did not correlate with *Cyp6g1* expression. A number of these metabolites produced in the *D. melanogaster*, such as IMI-5-OH, IMI-Ole and IMI-NNO, are known to be toxic to insects^36-38^.

Here the IMI metabolites produced by *D. melanogaster* and the subset of these formed by CYP6G1 are identified. Evidence is presented for the existence of other fly enzymes capable of producing these same metabolites. Rapid excretion of toxic metabolites produced by CYP6G1 in either the metabolic tissues or the CNS is observed. Indeed, we have not found evidence of IMI detoxification in *D. melanogaster*, underlining the significance of the transport and excretion of metabolites after their production in the insect. Lastly, we observed the presence of IMI-5-OH and IMI-Ole metabolites in a *Cyp6g1* knock-out strain, confirming the presence of at least one other metabolic gene capable of producing these IMI metabolites in this insect.

## Results

### Microbial degradation of IMI

Nitro-reduction of IMI is a biotransformation mechanism commonly found in soil bacteria^26,27^. In testing the hypothesis that the IMI-NH and IMI-NNO metabolites previously observed in *D. melanogaster*^35^ are produced by gut microbes, we examined the levels of these metabolites in axenic and non-axenic control larvae.

Neither IMI-NNO nor IMI-NH was detectable in axenic or control larvae exposed to IMI, but these metabolites were detected in the media in which both types of larvae were exposed. Significantly, the levels were 5 times lower in the media in which axenic larvae had been exposed (Figure 2, A). In contrast, there were no significant differences between the levels of the oxidative metabolites, IMI-5-OH and IMI-Ole, detected in the axenic and control larvae. Indeed, the levels of IMI-5-OH in the axenic larvae and their control were 2.4 ±1.1 and 2.8 ±0.2 ppb while the levels of IMI-Ole were 9.4 ±5.4 and 7.4 ±0.6 ppb respectively. Similar observations were made comparing the levels of these metabolites in the media in which the axenic and control larvae were exposed (Figure 2, B and C). IMI-diol and IMI-de metabolites were not detected at these experimental conditions.

**Figure 2.**
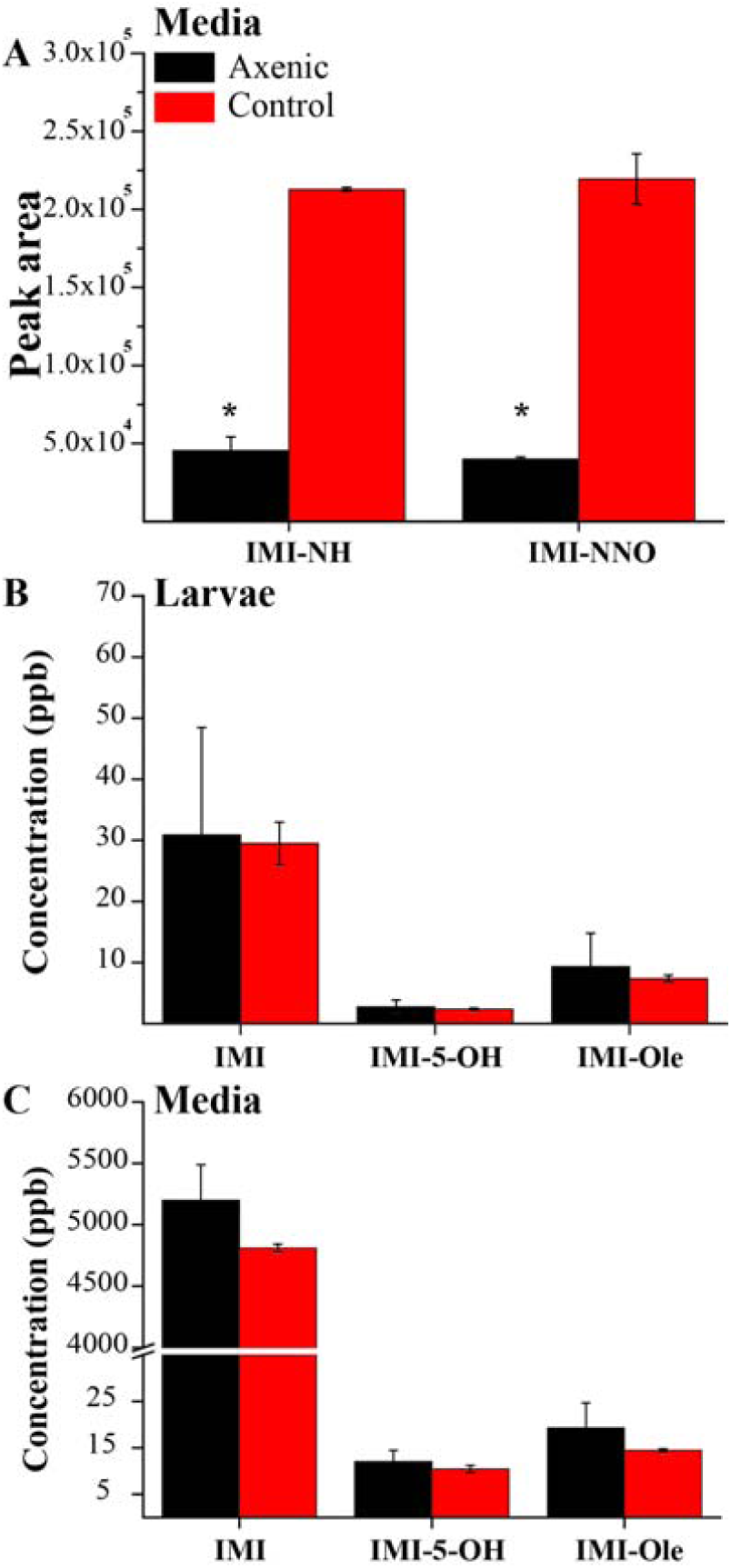
Metabolism in axenic larvae. (A) The amount of IMI-NH and IMI-NNO (reported as peak area) retrieved in the exposure media for axenic larvae (black) and controls (red). (B-C) The concentration of IMI, IMI-5-OH and IMI-Ole retrieved from the larvae and the exposure media for axenic and control larvae after a 6 hr exposure in a 6ppm solution of 12C_6_ -^13^C_6_ IMI (50:50 ratio). Significance is indicated by an asterisk and is calculated by comparing the levels of each metabolite detected in the axenic strain compared with the control. [The data represent the mean±SD; (n=3); Student-t-test: P≤0.01 (**), P≤0.05 (*)].

### The metabolites produced by CYP6G1

Previous studies have shown that *in vivo* over-expression of *Cyp6g1* in metabolic tissues of *D. melanogaster* using the GAL4/UAS system leads to increased production and excretion of IMI-5-OH, IMI-Ole, IMI-diol and IMI-de metabolites^35^, thus associating the action of CYP6G1 to the metabolism of IMI. However, it is still unknown which of these reactions are actually catalysed by CYP6G1 *in vivo*. In order to clarify the pathway of IMI metabolism and the steps catalysed by CYP6G1, HR_Cyp6g1 and control larvae were exposed to IMI and IMI-5-OH and the oxidative metabolites produced in the larvae and excreted in the media monitored using LC-MS.

Following exposure to IMI for 6 hrs, two times less IMI was detected in the bodies of HR_Cyp6g1 larvae compared to the control. Levels of the metabolites IMI-5-OH and IMI-Ole were 1.7 and 3.7 times higher in the HR_ Cyp6g1 larvae at this time. IMI-Ole was the most abundant metabolite produced, reaching the concentration of 118 ±5.7 and 32 ±2.2 ppb in the HR_Cyp6g1 and HR_Φ86FB larvae, respectively (Figure 3, A). IMI-5-OH and IMI-Ole concentrations were higher in the media in which larvae had been exposed, with significantly more metabolites excreted by the HR_Cyp6g1 strain. 7.8 times more IMI-5-OH and 11.5 times more IMI-Ole were excreted by HR_Cyp6g1 compared to the control strain (Figure 3 B). The IMI-diol (Figure 3, C) and IMI-de (Supplementary Figure S1) metabolites were detected only at low levels in the media. Their detection was only possible due to the presence of the ^13^C-isotope, which permitted the metabolite to be distinguished from the background noise.

**Figure 3.**
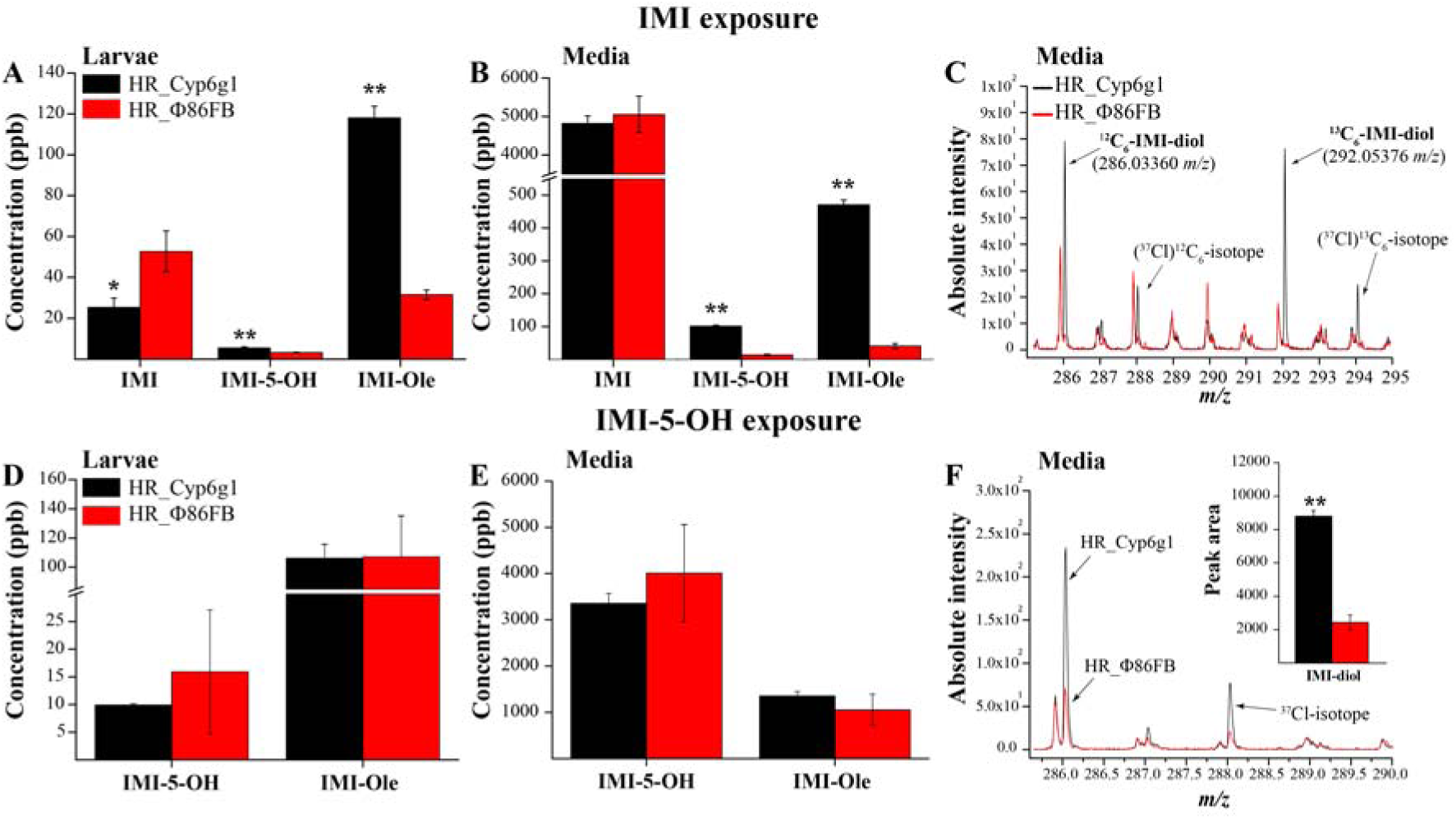
The metabolism of IMI and IMI-5-OH reported in the larvae and media of the HR_Cyp6g1 (black) and HR_Φ86FB (red) strains. Significantly less IMI and significantly more IMI**-**5**-**OH and IMI-Ole are detected in the HR_Cyp6g1 larvae and their respective media after exposure to IMI for six hrs. More IMI-Ole than IMI-5-OH was detected in these conditions (A-B). The IMI-diol metabolite (286.03360 *m/z*) was detected only in the HR_Cyp6g1 media. Its identification was facilitated by to the isotopic mass difference of ≈6.0201 mass units between the ^12^C_6_ and the ^13^C_6_-metabolites and the presence of the chlorine isotope (^37^Cl) (C). Exposure to IMI-5-OH revealed that HR_Cyp6g1and HR_Φ86FB strains produce and excrete equal levels of IMI-Ole (D-E). Significantly higher levels of IMI-diol were detected in the HR_Cyp6g1 media (black line) compared to the control (red line) (F). Significance (indicated with asterisks) is determined comparing the levels of the metabolites produced and excreted by the HR_Cyp6g1 strain against its HR_Φ86FB control [The data represent the mean±SD; (n=3); Student-t-test: P≤0.01 (**), P≤0.05 (*)].

The same genotypes were exposed to IMI-5-OH and the larvae and media were again analysed for metabolites (Figure 3 D-F). Only IMI-diol and IMI-Ole were detected. IMI-diol was only detected in the media where the levels were 3.6 times higher for the HR_Cyp6g1 genotype than the control (Figure 3 F). IMI-Ole was detected in the HR_Cyp6g1 and the control larvae (107 ±9.7 and 106±28.2 ppb) and in their respective exposure media (1300 ±97.5 and 1100 ±340.7 ppb). Differences in IMI-Ole levels between genotypes observed were not significant, suggesting that CYP6G1 did not catalyse the formation of this metabolite (Figure 3 D and E). Exposure to IMI-Ole did not lead to the detection of any other previously identified metabolites (Supplementary Figure S2).

### The kinetics of IMI metabolism when *Cyp6g1* is over-expressed in metabolic tissues

The over-expression of *Cyp6g1* in metabolic tissues has been previously shown to confer IMI resistance, as measured by the Wiggle index (WI) assay^39^. In that study a significant difference in the level of movement between the HR_Cyp6g1 and control HR_Φ86FB larvae was observed after 15 mins of IMI exposure. Here we chose to follow the kinetics of IMI metabolism in these same genotypes over a two hr time course, monitoring the levels of IMI metabolites produced and excreted over time.

Consistent with the WI data, overexpression of *Cyp6g1* in metabolic tissues leads to increased IMI metabolism (Figure 4). After 60 mins of exposure, the level of IMI present within the larval body was significantly lower for the HR_Cyp6g1 larvae. The levels of IMI in the media did not show any significant difference over time. (Figure 4 A and B).

**Figure 4.**
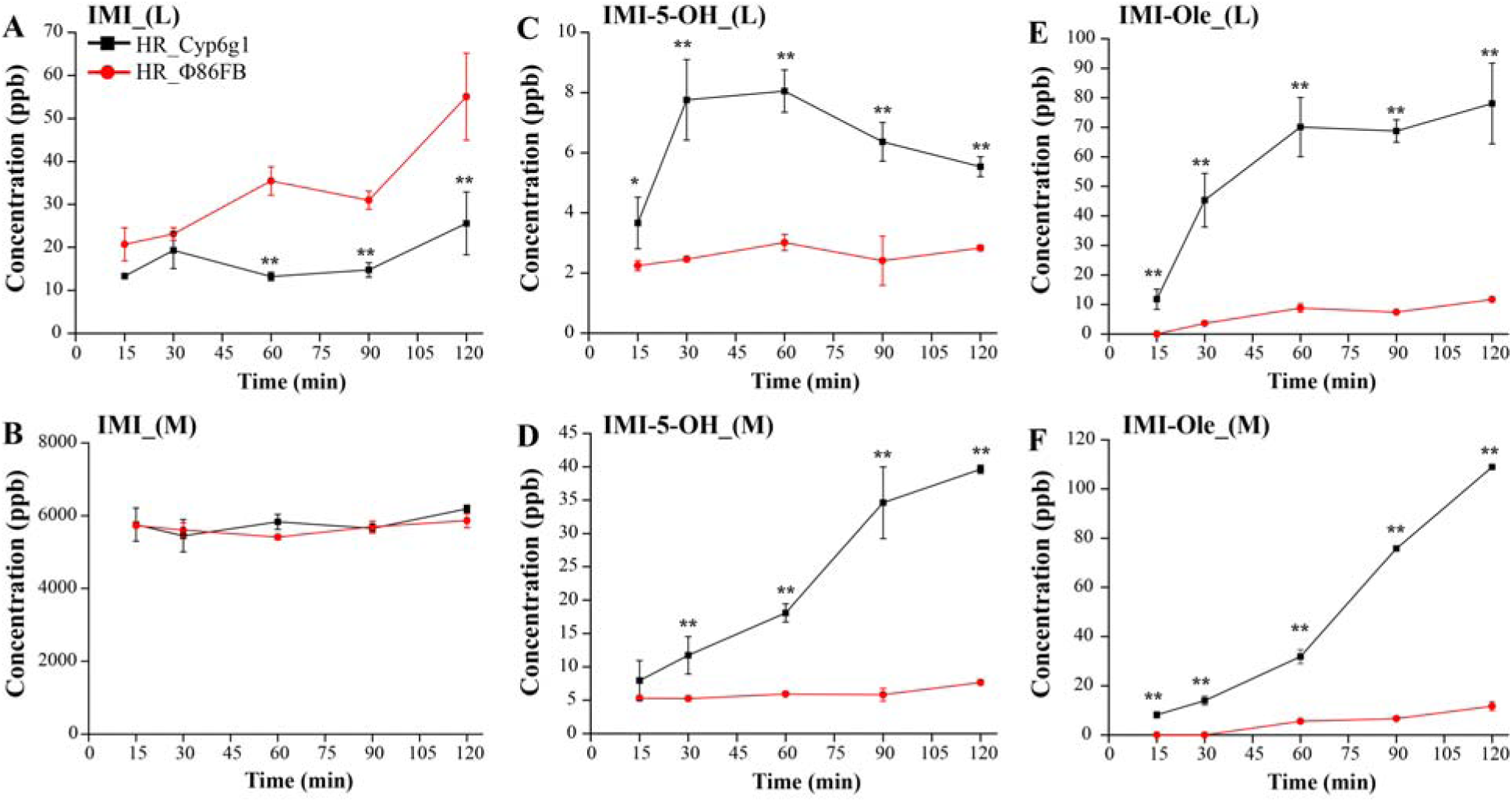
Time course metabolism of IMI in the HR_Cyp6g1 (black) and HR_Φ86FB (red) strains. The profile of IMI (A-B), IMI-5-OH (C-D) and IMI-Ole (E-F) detected in larvae (L) and media (M) over two hr assay. Significance (indicated with asterisks) is determined comparing the levels of the metabolites produced and excreted by the HR_Cyp6g1 strain against its HR_Φ86FB control [The data represent the mean±SD; (n=3); Student-t-test: P≤0.01 (**), P≤0.05 (*)].

A sharp increase in IMI-5-OH levels was observed after 15 mins in the HR_Cyp6g1 larvae. The maximum concentration of IMI-5-OH was reached between 30 and 60 mins (8 ±0.7 ppb). This was followed by a steady decrease in the second hr of exposure. In contrast, levels of IMI-5-OH did not change in the control larvae over the two hr time course. Excreted IMI-5-OH was observed in the media for both genotypes at the 15 min time point. After 30 mins, significantly higher levels of IMI-5-OH were detected in the media for the HR_Cyp6g1 genotype compared to the control. After 2 hrs, 5 times more IMI-5-OH had accumulated in the media for the HR_Cyp6g1 genotype compared to the control (Figure 4, C and D).

IMI-Ole was present at significantly higher levels in the bodies of HR_Cyp6g1 larvae compared to control at all time points. In the HR_Cyp6g1 larvae, a sharp increase in the levels of IMI-Ole in the first 60 mins was followed by a small increase in the next 60 mins. For the control larvae, IMI-Ole was not detectable at 15 mins. IMI-Ole then gradually increased to a final concentration of 11.7 ±0.4 after 2 hrs, 7 fold lower than what was observed in the HR_Cyp6g1 strain. In examining the media, significantly more IMI-Ole was excreted by the HR_Cyp6g1 larvae compared throughout the time course. The levels of IMI-Ole excreted by the HR_Cyp6g1 genotype, first detected at 15 mins, gradually increased over the first 60 mins. The rate of excretion increased in the second half of the experiment. For the control, IMI-Ole was not found in the media until 60 mins with a slight increase occurring in the second hr. After two hrs, 10 times more IMI-Ole was detected in the media of the HR_Cyp6g1 genotype compared to the control (Figure 4 E and F).

### The over-expression of *Cyp6g1* in the Central nervous system (CNS) provides an insight into the transport of the IMI-5-OH and IMI-Ole metabolites in larvae

Over-expression of *Cyp6g1* in the CNS has been shown to significantly reduce the impact of IMI on larval movement after 30 mins of exposure^39^. These data suggest that CYP6G1 can reduce the effect of IMI in the CNS by quickly metabolising it. To test this hypothesis, we examined the levels of IMI and metabolites over a two hr time course in the Elav_Cyp6g1 and Elav_Φ86FB control larvae.

Similar to the observations in the HR_Cyp6g1 experiment, IMI was rapidly metabolised into IMI-5-OH and IMI-Ole in Elav_Cyp6g1 larvae (Figure 5). However, the levels of IMI detected in the Elav_Cyp6g1 larvae were significantly different to the control only at the 2 hr time point (P<0.05). The amount of IMI in the media was equal for the two genotypes at all times. (Figure 5 A and B).

**Figure 5.**
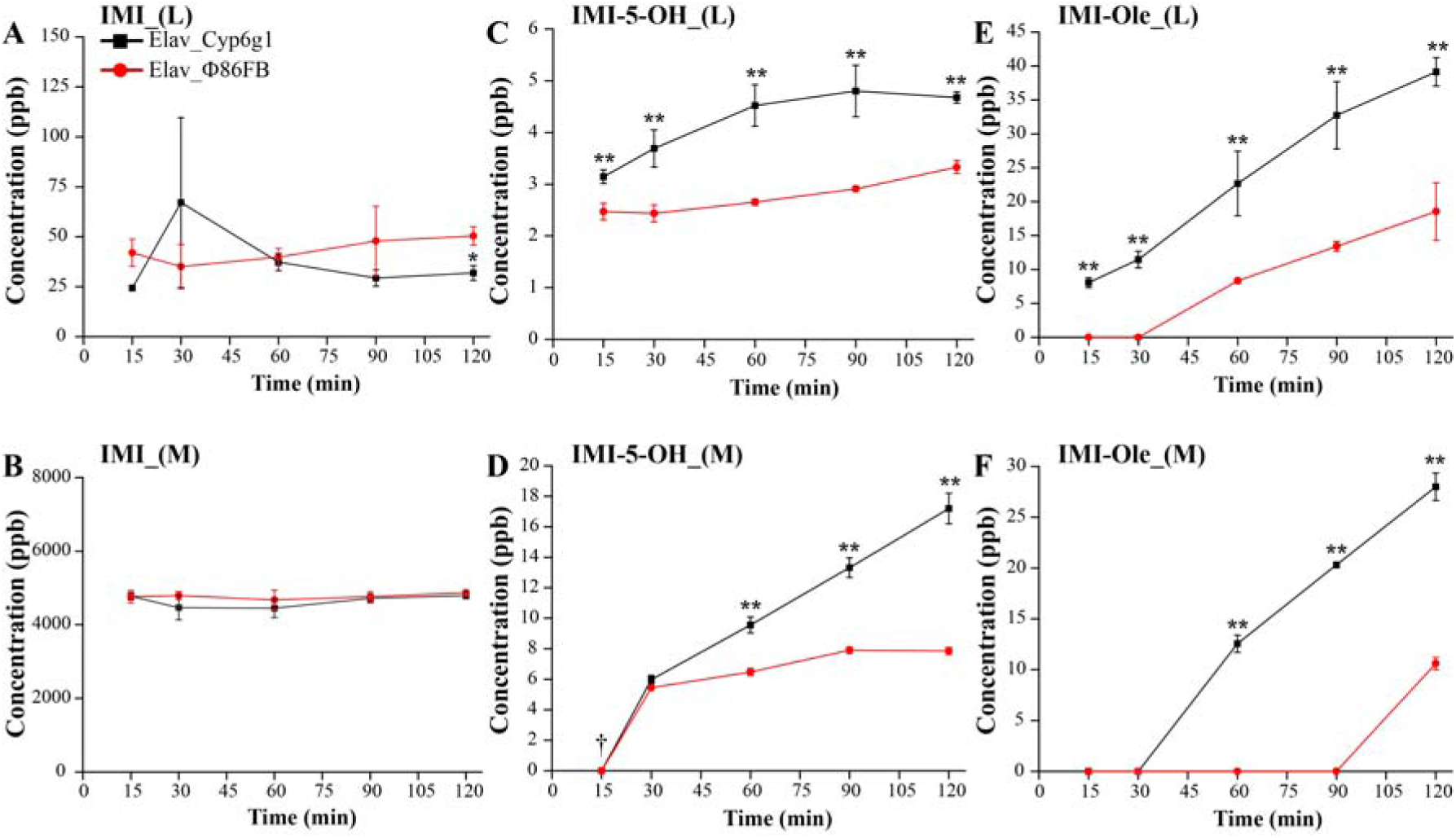
Time course metabolism of IMI in the Elav_Cyp6g1 (black) and Elav_Φ86FB (red) strains. The profile of IMI (A-B), IMI-5-OH (C-D) and IMI-Ole (E-F) detected in larvae (L) and media (M) over two hr exposure period. IMI-5-OH was successfully detected in the media of both genotypes at 15 minutes (†) but its intensity was below the LOQ. Significance (indicated with asterisks) is determined comparing the levels of the metabolites produced and excreted by the Elav_Cyp6g1 strain against its Elav_Φ86FB control [The data represent the mean±SD; (n=3); Student-t-test: P≤0.01 (**), P≤0.05 (*)].

IMI-5-OH was identified in the larval bodies of both strains after 15 mins of IMI exposure. Consistently higher levels of IMI-5-OH were detected in the Elav_Cyp6g1 larvae with 1.4 times more metabolite present after 2 hrs compared to the Elav_Φ86FB control. IMI-5-OH was identified in the media of both strains starting from 15 mins. After 60 mins significantly more IMI-5-OH was excreted by the Elav_Cyp6g1 genotype. Over the two hr assay, 2.2 times more IMI-5-OH was excreted by the Elav_Cyp6g1 genotype compared to the Elav_Φ86FB control (Figure 5 C and D).

IMI-Ole was rapidly formed in the Elav_Cyp6g1 larvae being detected at the 15 min time point while this metabolite was first detected in the Elav_Φ86FB controls starting from 60 mins. Over the two hr exposure, 2.2 times more IMI-Ole was produced by Elav_Cyp6g1 larvae compared to the Elav_Φ86FB control. IMI-Ole was detected in the media of the Elav_Cyp6g1 genotype starting from 60 mins while one more hr was required to observe IMI-Ole in the media for the control. After 2 hrs, 2.6 times more IMI-Ole was excreted by Elav_Cyp6g1 larvae (Figure 5, E and F).

### Measuring IMI metabolism in a CYP6G1 knock-out strain

Many other DMEs are expressed in the metabolic tissues of *D. melanogaster* larvae^39^ but their capacity to contribute to IMI metabolism and resistance remains largely untested. Denecke *et al.* (this issue) presented evidence that the selective knock-out of the two copies of the *Cyp6g1* gene from the insecticide resistant strain RAL_517 (RAL_517-Cyp6g1KO) significantly increases the susceptibility to IMI. As this would imply a reduction or loss of IMI metabolism in the insect, we verified this hypothesis analysing the change in metabolism of IMI in the larvae and media of the RAL_517 and RAL_517-Cyp6g1KO strains respectively.

Under the experimental conditions used in this research, loss of *Cyp6g1* function affected the metabolism of IMI in the RAL_517-Cyp6g1KO strain compared to the RAL_517 control. 1.5 and 3.8 times less IMI-5-OH and IMI-Ole, respectively, were detected in the RAL_517-Cyp6g1KO larvae compared to the RAL_517 control (Figure 6A). As a consequence, 1.8 times more IMI was detected in the body of the RAL_517-Cyp6g1KO larvae. RAL_517-Cyp6g1KO also excreted less IMI-5-OH and IMI-Ole than the RAL_517 control. 10 and 12.8 times less IMI-5-OH and IMI-Ole were detected in the media of RAL_517-Cyp6g1KO compared to the RAL_517 control (Figure 6B). Low levels of IMI-diol and IMI-de metabolites were detected only in the media of the RAL_517 genotype. A similar metabolic response was observed at the same experimental conditions reported by Denecke *et al* (this issues) (Supplementary Figure S4). Although the loss of CYP6G1 in *D. melanogaster* drastically affected the metabolism of IMI, the oxidative metabolites, IMI-5-OH and IMI-Ole, were still observed in the larval bodies and exposure media of the RAL_517-Cyp6g1KO strain at both exposure conditions used. Therefore CYP6G1 cannot be considered the only enzyme responsible for the metabolism of IMI in the insect.

**Figure 6.**
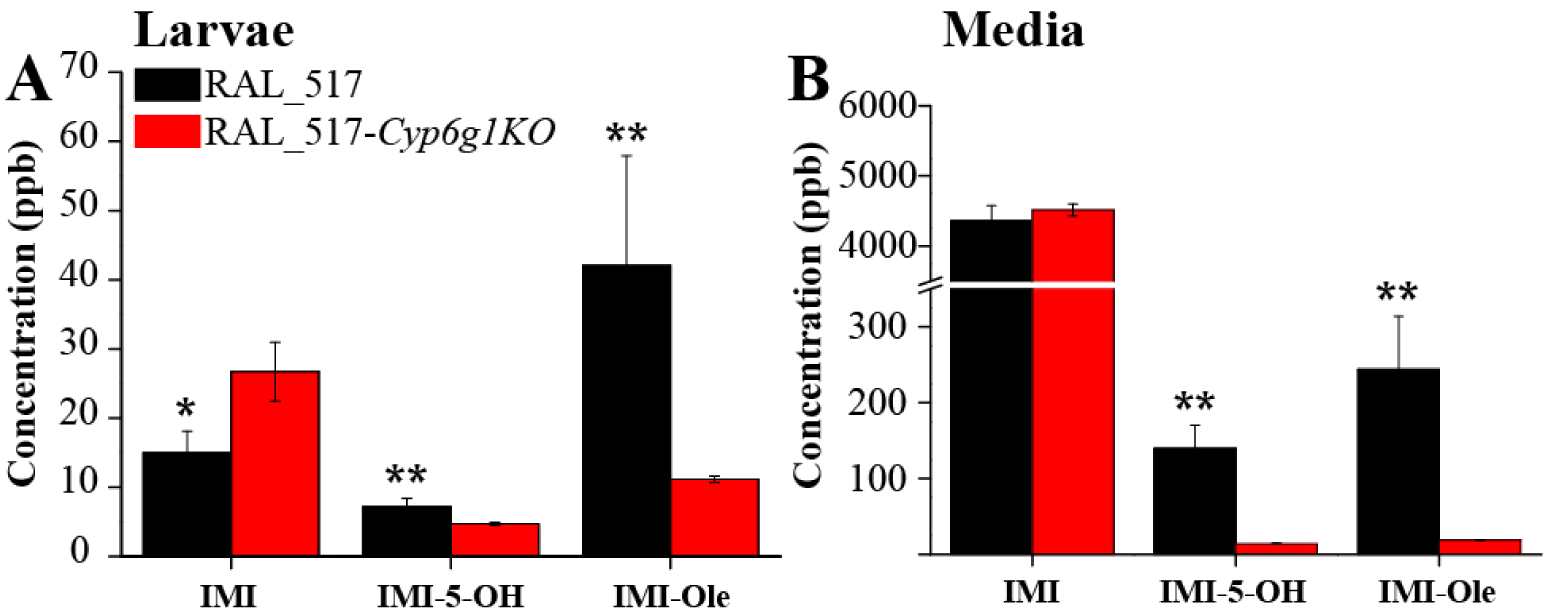
The metabolism of IMI detected in the larvae (A) and media (B) of the RAL_517 (black) and RAL_517-Cyp6g1KO (red) strains. Metabolism of IMI was identified after 6 hrs of exposure to a 6 ppm solution of IMI. Significantly more IMI-5-OH and IMI-Ole are produced and excreted by RAL_517 compared to the knock-out strain. Significance (indicated with asterisks) is determined comparing the levels of the metabolites produced and excreted by the RAL_517-Cyp6g1KO genotype with the RAL_517 parental control [The data represent the mean±SD; (n=4); Student-t-test: P≤0.01 (**), P≤0.05 (*)].

## Discussion

Like many other insecticides, the targets of IMI are in the CNS. Hence, toxicity depends on the concentrations of insecticide and/or toxic metabolites that enter the CNS and the extent to which they can be effluxed from it. The major metabolites of IMI produced in *D. melanogaster* had been previously defined. These included the oxidative metabolites IMI-5-OH, IMI-Ole and IMI-diol, the de-ethylation product of IMI, IMI-de, the nitro-reduced derivatives IMI-NNO and IMI-NH, the epoxidation product of IMI and the hydroxyl-derivative of IMI-NH^35^. The latter two were not detected in our study. Using tissue specific gene expression our data allows the *in vivo* contribution of one gene, *Cyp6g1*, to IMI metabolism to be characterised and quantified. CYP6G1 is revealed to be an enzyme that produces metabolites that are toxic but readily excreted, underlining the significance of yet to be identified transporters in the response to IMI. As shown by our data, metabolites can be partitioned into those produced by *D. melanogaster* (oxidative metabolites) and those produced by microbes within the fly (nitro-reduced metabolites).

Microbes living within the *w*^*1118*^ strain make a significant contribution to the nitro-reduction of IMI (Figure 2). In the absence of microbes the amount of IMI-NNO and IMI-NH excreted by the axenic *w*^*1118*^ larvae was significantly reduced, while oxidative metabolism was unaffected. In Aphids, IMI-NNO is more toxic than IMI, while IMI-NH has low level toxicity^38^. While the toxicity of these compounds has not been tested here, it appears likely that they do not reach the CNS, as they were not detected in larval bodies but were present in the exposure media (Figure 2B). IMI-NNO and IMI-NH metabolites can therefore be defined as microbial metabolites (Figure 7). The identification of IMI-NH and IMI-Ole-NH metabolites (Figure 1) in *R. flavipes*^33^ and the identification of IMI-Urea in metabolic tissues of *A. mellifera*^*32*^ reveals that nitro-reduction of IMI is also present in other insects. Although the role of gut bacteria in resistance to different plant secondary metabolites (e.g. terpenes and glucosinolates) have been reported in several species^40-42^, evidence of their possible contribution in insecticide resistance^43^ and metabolism^44^ is more limited. The role of microbiota in resistance to insecticides is still at its infancy and more studies are required to address this question in detail. Nonetheless, in future caution must be applied in attributing levels of insecticide metabolism and resistance to the over-expression of insect metabolic enzymes without first investigating whether there is a microbial contribution.

**Figure 7.**
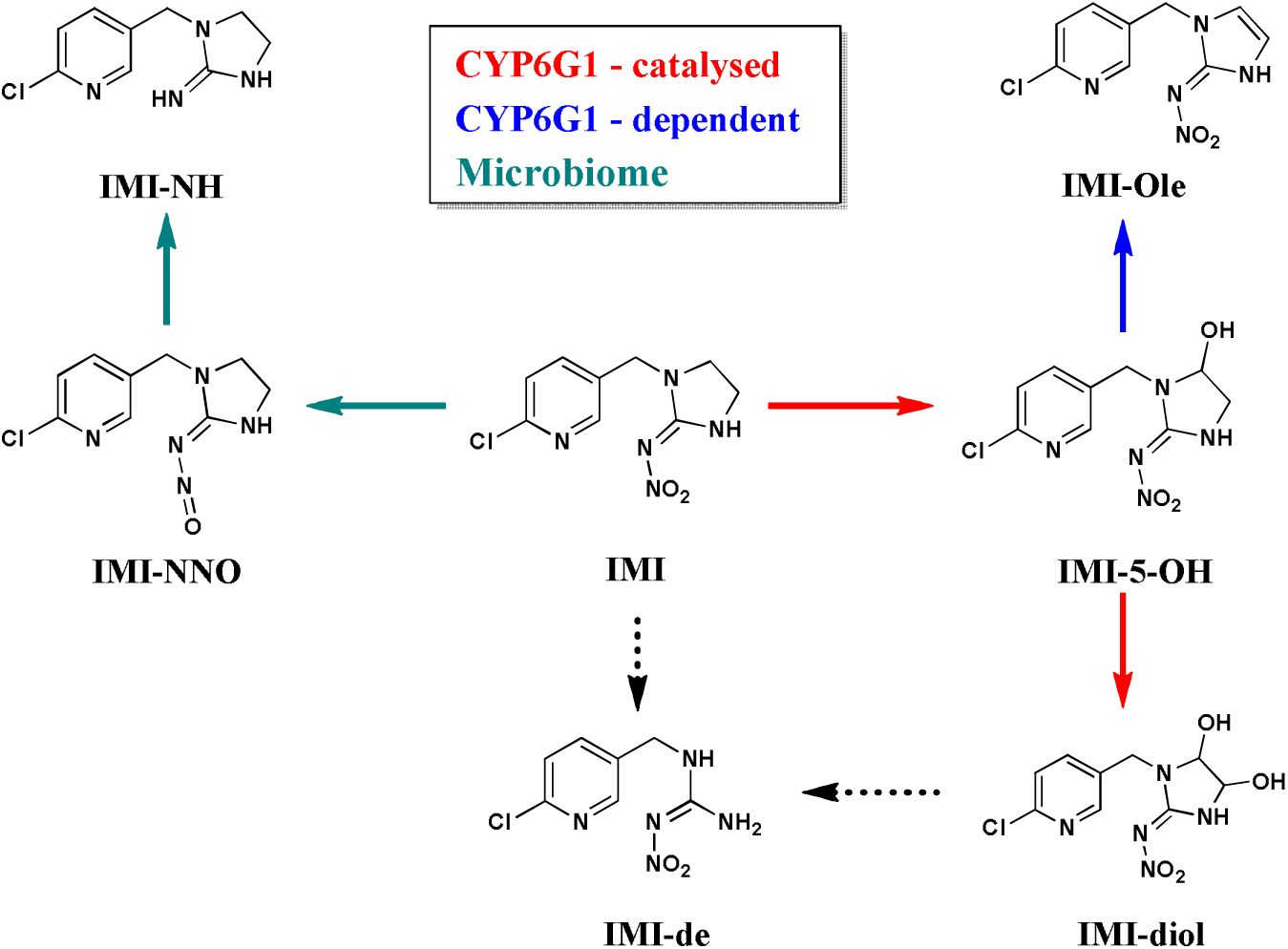
The metabolism of IMI in *D. melanogaster.* Red, blue and green arrows represent CYP6G1-catalysed, CYP6G1-dependent and microbial metabolites respectively. Dashed arrows leading to IMI-de are used to indicate the two possible pathways by which IMI-de could be formed.

*D. melanogaster* typically harbour a limited number of bacterial species in the gut that are capable of establishing a strong symbiotic relationship with the host by regulating different physiological mechanisms included growth, immune response and the recycling of nutrients^45^. Bacteria within the larval midgut would be exposed to orally ingested IMI, as would the *D. melanogaster* metabolic enzymes expressed there^46^. Bacteria with the capacity to metabolise insecticides have been used as tools for the bioremediation of insecticide residues in the environment^26,47^. Hence, there may be value in identifying the species responsible for the IMI metabolism reported here. Many of the bacterial species commonly found in *D. melanogaster* can be cultured, so this may be possible.

That the levels of IMI-NNO and IMI-NH were not reduced to zero in axenic larvae is consistent with the hypothesis that there are one or more *D. melanogaster* enzymes that catalyse the nitro-reduction of IMI. CYP6G1 has been ruled out as a candidate for this activity^35^, since the over-expression of the *Cyp6g1* gene does not impact the levels of these metabolites. In looking for other enzymes capable of producing these metabolites, it is necessary to look beyond insects, since there are no reported examples of insect enzymes with this capacity. P450s in human^23^ and Alcohol Oxidase (AOX) in mouse^48^ have been shown to posses the ability to nitro-reduce IMI.

The oxidative metabolism of IMI in *D. melanogaster* is a complex pathway composed of several steps (Figure 7). We distinguish between two classes of oxidative metabolites, those formed by CYP6G1 activity (CYP6G1-catalysed) and those dependent on the expression of CYP6G1 (CYP6G1-dependent).

IMI-5-OH and IMI-diol are CYP6G1-catalysed metabolites. Oxidation of IMI in position 5 of the imidazolidine ring forms IMI-5-OH and then further hydroxylation in position 4 forms IMI-diol. This two-step process previously observed *in vitro*^34^ has been shown here to occur *in vivo*. Exposure to IMI-5-OH revealed a significant increase in the IMI-diol detected in the HR_Cyp6g1 media compared to the control (Figure 3F). Jouβen, *et al* (2008)^34^ identified IMI-4-OH as a CYP6G1 catalysed metabolite. We have not been able to detect this metabolite in *D. melanogaster*.

In mice and plants, IMI-Ole is formed by dehydration of IMI-5-OH^22,24,38^. *In vitro*, studies indicated that CYP6G1 does not catalyse the conversion of IMI-5-OH into IMI-Ole^34^. This was confirmed to be the case *in vivo* by the exposure of the HR_Cyp6g1 and control genotypes to IMI-5-OH (Figure 3D and E). The over-expression of *Cyp6g1* in the metabolic tissues of the HR_Cyp6g1 genotype does not affect levels of IMI-Ole produced and excreted by this strain compared to the control. As such, it can be deduced that the increasing levels of IMI-Ole observed in the HR_Cyp6g1 strain when exposed to IMI are due to the increasing amount of the substrate, IMI-5-OH, produced by the over-expression of *Cyp6g1* in metabolic tissues (Figure 3A and B).

The IMI-de metabolite observed in spinach^24^, and previously detected in *D. melanogaster*^35^, has also been observed here. IMI-de formation can be explained by de-ethylation on the imidazolidine ring of IMI^24^ or loss of ethylene glycol from the IMI-diol metabolite. Detection of IMI-de in the media of the HR_Cyp6g1 strain after exposure to IMI (Supplementary Figure S1), but not after exposure to IMI-5-OH, indicates that IMI-de may be produced by IMI de-ethylation. That it was not detected in the HR_Φ86FB strain suggests that CYP6G1 could play a role in this conversion.

By quantifying IMI metabolites, we established that over-expression of *Cyp6g1* in the metabolic tissues increases the amount of the oxidative metabolites within larvae and the amount excreted (Figure 3A and B). This observation was confirmed in the two hr time course assay where only IMI-5-OH and IMI-Ole were detected (Figure 4). Significantly higher levels of metabolites detected in the HR_Cyp6g1 larvae and their media between 15 and 30 mins suggest that ingested IMI is rapidly metabolised by CYP6G1 in the metabolic tissues and that IMI-5-OH and IMI-Ole are excreted in the media of exposure as they are formed. These data are consistent with a previous report that there is a significant difference in motility between the HR_Cyp6g1 and control strains after only 15 mins of exposure to IMI^49^. The over-expression of *Cyp6g1* in the metabolic tissues reduces the levels of IMI in larval bodies reducing the toxicity of this insecticide (Figure 4A).

The 60 min time point marks a noteworthy change in the trends observed for IMI-5-OH and IMI-Ole levels in the HR_Cyp6g1 strain (Figure 4C-F). Rapid metabolism of IMI in the first 60 mins of exposure, observed as increase of IMI-5-OH and IMI-Ole, is accompanied by a proportional amount of these metabolites being released into the media. After 60 mins, the production of IMI-5-OH and IMI-Ole slows down in the body with not sensitive change while excretion continues at a high efficiency. This process is visible after 6 hrs of exposure of the HR_Cyp6g1 strain to either IMI or IMI-5-OH. While similar levels of IMI-Ole can be detected for both chemical used, 4 and 13 times more IMI-Ole is excreted by this strain upon exposure to IMI and IMI-5-OH respectively (Figure 3).

A similar metabolic trend is observed with the over-expression of *Cyp6g1* in the CNS (Figure 5). Significant levels of IMI-5-OH and IMI-Ole observed after 15 mins in the Elav_Cyp6g1 larvae indicate that IMI can reach the CNS in a relatively short time where it is metabolised by CYP6G1 into IMI-5-OH. While the detection of IMI-5-OH in the media in the first 30 mins reflects the base-level expression of *Cyp6g1* in the metabolic tissues of both strains, the significantly higher levels of IMI-5-OH and IMI-Ole observed after 60 mins indicate that metabolites produced in the CNS are excreted out of the larvae. For this to occur, metabolites produced in the CNS must cross the blood brain barrier, diffuse into the hemolymph and reach the malpighian tubules for excretion into the media. As the production of IMI-Ole is not catalysed by CYP6G1, it is not clear whether it is produced in the CNS or not.

Although resistance is attributed to the over-expression of *Cyp6g1*, CYP6G1 does not actually detoxify IMI. It rather produces IMI-5-OH, which leads to the production of IMI-Ole. Both metabolites are toxic in *D. melanogaster* and are likely to bind to at least one nAChR subunit targeted by IMI (Supplementary Figure S3). That these metabolites are less toxic than IMI in *D. melanogaster* could be explained if they are less able to enter the CNS, have a lower binding affinity for nAChRs and/or if they are more readily excreted. That excretion plays a vital role in the capacity of an insect to survive to insecticide exposure is particularly evident with IMI-Ole, a metabolite produced at high concentration and known to be more toxic than IMI in pest species like *M. persicae*, *A. gossypii*^38^ and *Bemisia tabaci*^37^, and in the honeybee *Apis mellifera*^36^. While the basis of these differences in toxicity between species have not been established, they can be explained by mechanisms canvassed here. In the case of *D. melanogaster* our data show rapid excretion of IMI-5-OH, IMI-Ole, IMI-de, IMI-NNO and IMI-NH. The capacity to transport and excrete these metabolites is crucial to the survival under conditions of exposure to toxic concentrations of IMI. This has been reported by Denecke *et al* (this issue) where the hypersensitivity of one strain (RAL_509 strain) to IMI can be explained by its reduced capacity of eliminate the toxic metabolites IMI-5-OH and IMI-Ole, that accumulate within the insect body. CYP6G1 metabolises IMI, but detoxification depends on the excretion of IMI and the metabolites produces. Our data, combined with evidence linking candidate genes such as ABC transporters to insecticide resistance^50,51^, suggest that the sequential activity of DMEs and transporters may be important contributor to the resistance to a range of insecticides. More research needs to be devoted to identifying the specific transporters involved in moving insecticides/metabolites between tissues to the point of excretion.

Finally, the analysis of a *Cyp6g1* knock-out mutant revealed the presence of metabolites normally associated with CYP6G1 activity, indicating functional redundancy and confirming the role of at least one other gene in the metabolism of the insecticide. Our data show that the increased susceptibility to IMI observed with the RAL_517-Cyp6g1KO genotype (Denecke *et al.* this issue) correlates with a significant decrease in production of IMI-5-OH and IMI-Ole compared to the control RAL_517 (Figure 6). A similar metabolic result is observed after only one hour of exposure at the same conditions used by (this issue) (Supplementary Figure S4). However, the persistence of IMI-5-OH and IMI-Ole metabolites in the RAL_517-Cyp6g1KO strain, albeit at lower levels, indicates that one or more metabolic enzymes can contribute to the metabolism of IMI. That the same metabolites normally produced by *Cyp6g1* are detected suggest that a P450 may be involved in this enzymatic conversion, possibly one closely related to *Cyp6g1*^52^. *Cyp6g2* is one candidate. Denecke *et al.* (this issue) have shown that there is a significant level of expression of this gene in the metabolic tissues of RAL_517 and that transgenic overexpression of the gene leads to both elevated levels IMI-5-OH and IMI-Ole and IMI resistance. Any mutation that leads such a gene to be over-expressed in a similar tissue-specific pattern to *Cyp6g1* or in the CNS would be likely to confer insecticide resistance, in the absence of significant fitness costs. Candidate genes can be readily tested with controlled gene overexpression^21^ and their role in insecticides metabolism can be assessed using LC-MS.

## Materials and Methods

### Chemicals

IMI (N-[1-[(6-Chloro-3-pyridyl)methyl]-4,5-dihydroimidazol-2-yl]nitramide) and [^13^C_6_]-IMI (>99% isotopic purity; >97% total purity) were obtained from AK Scientific (Union City, CA, USA) and IsoSciences (King of Prussia, PA, USA), respectively. Authentic standards of IMI-Ole (N-[1-[(6-chloro-3-pyridyl)methyl]-1,3-dihydro-2H-imi-dazol-2-ylidene]nitramide) and IMI-5-OH (N-(1-[(6-chloro-3-pyridyl)methyl]-5-hydroxyimidazolidin-2-ylidene]nitramide) were provided by Bayer CropScience AG (Monheim, Germany). Sucrose was obtained from Chem-Supply (Gillman, SA, Australia) and glacial formic acid from AnalaR (supplied by BDH, Poole, Great Britain). HPLC grade acetonitrile, ethyl acetate, and HPLC grade methanol were obtained from Merck (Kenilworth, NJ, U.S.A.). 18.2 M□ HPLC grade water was obtained from Honeywell (Morristown, NJ, USA).

### *D. melanogaster* genotypes and rearing conditions

Tissue-specific over-expression of *Cyp6g1* was used to identify and monitor metabolites produced by CYP6G1. Over-expression was achieved by crossing the 5’HR_GAL4 Hikone R (Accord) driver line ^53^ or the Elav_GAL4 (<u>Bloomington Drosophila Stock Center</u> #458) driver line with a UAS-Cyp6g1 line^46^. These crosses generate progeny, which over-express *Cyp6g1* in metabolic tissues (UAS_Cyp6g1-86FB × 5’HR_GAL4, referred to as HR_Cyp6g1) or in the CNS (UAS_Cyp6g1-86FB × Elav_GAL4 referred to as Elav_Cyp6g1). A control genotype, differing only in *Cyp6g1* tissue-specific expression levels, was generated by crossing the 5’HR_GAL4 and Elav_GAL4 driver lines to the Φ86FB genotype^54^ referred to as HR_Φ86FB and Elav_Φ86FB respectively. Flies were reared at 25 °C and raised in bottles filled with rich media food (Supplementary, Table S1).

### Axenic larvae preparation

Axenic larvae of *D.melanogaster* were allowed to develop from embryos that were decontaminated and dechorionated under a laminar flow cabinet^55^. Flies from the susceptible *w*^*1118*^ strain (<u>Bloomington Drosophila Stock Center</u> #3605) were collected in cages and allowed to lay eggs on grape juice plates (Supplementary information, Table S1). Every 2 hrs, cages were moved onto new grape juice plates. Freshly laid embryos were collected and sterilized using a dilute solution of bleach (2.5% v/v) for 2 mins, followed by a 30 second wash in ethanol (70% v/v) and finally rinsed with sterilized distilled water for one min. Eggs deprived of the external shell were selected under the microscope and gently transferred on autoclaved rich media food containing 50 and 20 mg/L of ampicillin and chloramphenicol, respectively. The glass vials used for the exposure, the lysing matrix tubes, and the different solutions of sucrose used in the experiment were autoclaved before use.

### Insecticide exposure and metabolite extraction process

The exposure and extraction conditions employed to test and extract IMI and metabolites from larvae of *D. melanogaster* were described elsewhere^35^. Briefly, eggs were laid on rich media food for 24 hrs. After 4 days, larvae were extracted from the food using a 15% sucrose solution. Larvae were gently transferred onto a mesh, rinsed with more sucrose solution and counted under a microscope to ensure that only early third instar larvae were used in the experiments. For each experiment or time point, three biological replicates of 200 early third instar larvae were prepared. Larvae were exposed to a fresh solution of 50:50 ratio of ^12^C_6_ and ^13^C_6_-IMI (total concentration 6ppm) dissolved in 200μL of a 5% (w/v) sucrose solution. For exposures to IMI-5-OH and IMI-Ole, the solution was composed only of the ^12^C isotope. At the end of each experiment, the exposure media and larvae were separated and transferred into lysing matrix tubes containing 1.4 mm ceramic beads (MP biomedical, Santa Ana, California, USA). 1000 μL of distilled water was used to wash larvae prior to collection. The biological samples were milled using a Cryomil (Precellys 24, Bertin Technologies, Montigny-le-Bretonneux, France) operated at −10°C. IMI and its metabolites were then extracted using ethyl acetate. The extraction was repeated three times and the fractions combined in 2.0 mL tubes and evaporated to dryness under vacuum.

### LC/MS analysis

The analysis of IMI and metabolites using TIM has been previously reported^35^. Briefly, dried samples were resuspended in 250 μL of HPLC grade water and analysed using an Agilent 1100 HPLC autosampler system with a reverse-phase C18 column (2.6 μm, 3.0 × 100 mm, Kinetex XB-C18, Phenomenex, Inc., Torrance, California, USA) coupled to an Agilent 6520 Q-TOF mass spectrometer (Agilent Technologies, Inc., Santa Clara, CA, USA). The ionisation of IMI and its metabolites was achieved using a dual-nebulizer ESupplementary source with the following settings: capillary voltage, +3 kV; gas temperature (nitrogen), 300° C; dry gas, 12 L min^-1^; nebulizer, 50 psig. The flow rate was maintained at 0.3 mL/min and the total run was about 25 mins. Separation was achieved using a two solvent gradient consisting of (Phase 1) 100% H_2_O and (Phase 2) a 70:30:0.1% (v/v) mixtures of acetonitrile:H_2_O:formic acid, respectively. IMI and its metabolites were analysed in both ionisation modes. Only IMI and its major metabolites IMI-5-OH and IMI-Ole were quantified. This was achieved using negative ionisation mode and through external calibration curves generated by diluted solutions of pure standards (n=8) ranging from 1 ppb (part per billion) to 3 ppm (part per milion). The signal-to-noise ratio (S/N) equal to 3:1 was chosen as Level of Detection (LOD) while a S/N equal to 5:1 was used as Level of Quantification (LOQ) since quantification is determined from the sum of the areas produced by both isotopes. The LOQ for IMI, IMI-5-OH and IMI-Ole was ≈1-2 ppb. The levels of the metabolites were averaged among the three biological replications of each genotype and a Student’s t-test was performed to determine if significant difference existed between any two genotypes being compared. The standard deviation was plotted with average values for each metabolite to represent the variability among the three biological replicates. In the text mean±SD is shown for metabolites.

## Acknowledgements

The authors thank the University of Melbourne for the provision of the postgraduate and writing up scholarships to Roberto Fusetto and Shane Denecke and to all members of the O’Hair and Batterham research groups for their support. We thank Metabolomics Australia and the University of Melbourne Department of Chemistry for access to facilities and Sioe See Volaric for technical assistance with the analytical instrumentation.

## Author contributions

R.F and Ph.B: Writing the manuscript

R.F, S.D, T.P, R.O’H and Ph.B: Editing the manuscript

R.F, S.D and T.P: Laboratory Work

R.F, S.D, T.P: Analysis of data

Ph.B, R.O’H: Funding and overall project design

## Additional information

**Supplementary information** accompanies this paper.

